# The Pseudo-Torsional Space of RNA

**DOI:** 10.1101/2022.06.24.497007

**Authors:** Leandro Grille, Diego Gallego, Leonardo Darré, Gabriela da Rosa, Federica Battistini, Modesto Orozco, Pablo D. Dans

## Abstract

The characterization of the conformational landscape of the RNA backbone is rather complex due to the ability of RNA to assume a big variety of conformations. These backbone conformations can be depicted by pseudo-torsional angles linking RNA backbone atoms, from which Ramachandran-like plots can be built. We explored here different definitions of these pseudo-torsional angles, finding that the most accurate ones are the traditional η (eta) and θ (theta) angles, which represent the relative position of RNA backbone atoms P and C4’. We explore the distribution of η-θ in known experimental structures, comparing the pseudo-torsional space generated with structures determined exclusively by one experimental technique. We found that the complete picture only appears when combining data from different sources. The maps provide a quite comprehensive representation of the RNA accessible space, which can be used in RNA-structural prediction. Finally, our results highlight that protein interactions leads to significant changes in the population of the η-θ space, pointing towards the role of induced-fit mechanisms in protein-RNA recognition.

## INTRODUCTION

RNA molecules participate in almost any genomic process. The repertoire of RNA molecules is growing continuously: we know from decades that messenger RNAs (mRNA) transports the genomic information which is transformed into proteins with the help of ribosomal and transfer RNAs, but very recently many other functional RNAs have emerged. For example, long non-coding (lncRNA) and small interfering RNA (siRNA) are essential for the regulation of gene expression and gene activation/silencing, and some micro RNAs (miRNAs) are known to trigger mRNA degradation. Small nuclear RNA (snRNA) is critical for the function of the spliceosome, and ribozymes are RNA molecules capable of catalysing specific biochemical reactions. This astonishing diversity in function is due to RNA ability to assume a surprisingly variety of conformations, which structurally speaking places the RNA closer to proteins than to DNA.

At physiological conditions, unlike DNA that is mostly found as a regular double helix, RNA molecules can adopt several secondary motifs. These can interact within each other, like kissing-loops, or even fold, like in pseudo-knots and junctions, to form highly complex 3D structures. Although only less than 5% of the structures determined experimentally and deposited in the Protein Data Bank (PDB) contain RNA, they sample a large variety of those secondary structure motifs. Among the most studied, hairpins with 3 to 11 bases in the loop region can be found. Some of them have particular structural elements, such as U-turns and A-turns typical of tRNA, while others display a particular sequence preference, like the known GNRA and UNCG tetra-loops common in rRNA. Symmetric and asymmetric internal loops also encompass a significant number of important motifs, like K-turn, C-loop, docking-elbow, bulged-G, and twist-up motifs, which in most of the cases are composed by a combination of important structural elements like S-turns, base triplexes, cross-strand stacking, and extruded nucleotides. In addition to this rich conformational landscape, RNA molecules are often found in functionally specific protein-RNA complexes, i.e. the ribosome or the spliceosome, where the effect of the protein on the RNA conformation is unclear.

The first attempt to rationalize and classify RNA motifs was based on the nucleobase pairing capabilities, i.e. the ability to form hydrogen bonds. Saenger’s classification (1) included 29 different base pairs in RNA. This classification was reduced a posteriori to 12 discrete groups by Leontis & Westhof (2), who also introduced a general nomenclature to describe base pairing rules. These groups describe canonical and non-canonical pairs, regardless of the structural motif. However, no information is given about the sugar-backbone conformation. Initial attempts to classify the RNA backbone conformational space led to 37 (3, 4) and 32 (5) backbone rotamers, highlighting the complexity of the RNA backbone when all the torsions are considered. In 2008, the RNA Ontology Consortium (ROC) updated and refined the clustering of the multidimensional dihedral angle distributions, into 46 discrete backbone conformers. Using a different approach, first Olson (6), and later on Pyle’s group (7), proposed the usage of two pseudo-bonds (traced between the C4’ and P atoms of the backbone), to simplify the backbone conformational space. Considering these pseudo-bonds, the dihedral space of the phosphodiester linkage can be described by only two pseudo-torsional angles: η (eta: C4’_i-1_, P_i_, C4’_i_, P_i+1_) and θ (theta: P_i_, C4’_i_, P_i+1_, C4’_i+1_) (Figure 1). “Ramachandran-like” plots of η-θ allowed to identify 11 major backbone conformers, discriminating according to the sugar conformation (North or South). It was demonstrated that nucleotides with similar η-θ values had almost identical backbone conformations, and a relationship was detected between the backbone conformation and the base position, pointing out the ability of η-θ to discriminate between base orientations.

**Figure 1.**
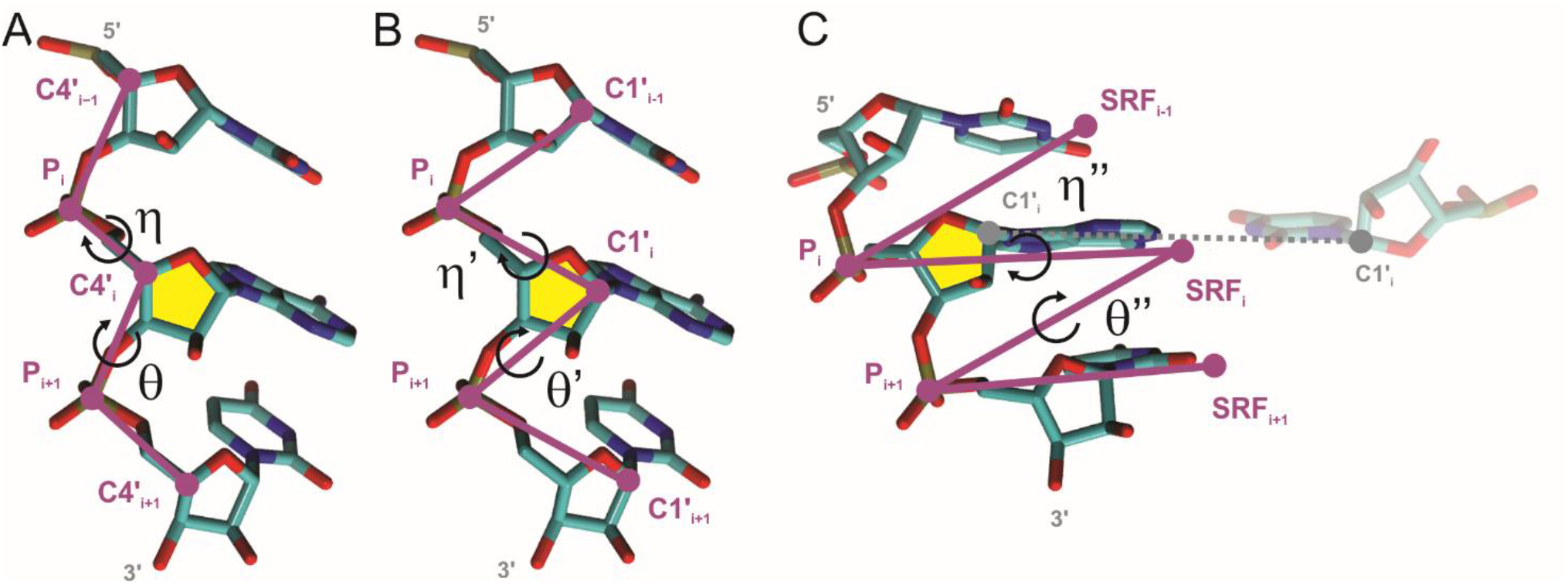
Structural representation of the pseudo-torsional angles of RNA. A) Definition of the η (eta: C4’_i-1_, P_i_, C4’_i_, P_i+1_) and the θ (theta: P_i_, C4’_i_, P_i+1_, C4’_i+1_) angles. B) Definition of the η’ (eta’: C1’_i-1_, P_i_, C1’_i_, P_i+1_) and the θ’ (theta’: P_i_, C1’_i_, P_i+1_, C1’_i+1_) angles. C) Definition of the η’’ (eta’’: SRF_i-1_, P_i_, SRF_i_, P_i+1_) and the θ’’ (theta’’: P_i_, SRF_i_, P_i+1_, SRF_i+1_) angles, where SRF stands for Standard Reference Frame as implemented in DSSR (9).

Aiming to increase structural characterization two other pseudo-torsional angles have been defined, namely η’-θ’, where the C4’ in η-θ was replaced by C1’ (8), and η’’-θ’’, formed between the phosphate atoms and a standardized point in the nucleobase plane (9), according with the standard reference frame (SRF) as defined by Olson and co-workers (10). η’-θ’ was less explored even though in principle it seems more adequate than η-θ to fit the RNA backbone to X-ray electron density maps. The η’’-θ’’ has the additional advantage that it contains information about the nucleobase orientation respect to the RNA backbone, something not explicitly provided by the other two pairs of pseudo-torsions. No comprehensive study has been published comparing the informational load of these pseudo-torsional angles.

In the last years, the number of structures determined by X-ray crystallography (X-ray), Nuclear Magnetic Resonance (NMR), or Cryogenic Electron Microscopy (Cryo-EM) techniques, and available in public databases, has grown to the point where there is enough, non-redundant, data to extract statistical meaningful information. This allows us to compare isolated RNA molecules *vs*. those bound to proteins, finding any potential bias related to the experimental sources of 3D structures, or to start adding sequence context to major conformers. Accordingly, in this work we used an updated version of our veriNA3d R (11) package to revisit the 3 pairs of pseudo-torsional angles (η-θ, η’-θ’, and η’’-θ’’) as they emerge from non-redundant experimental structures available to date. We found that η-θ were the best pseudo-torsions to describe RNA conformational space. We find 14 major states and detected non negligible differences in sampling of the different conformations depending on the technique used to determine the structure. Finally, the presence of binding can lead to the exploration of typically empty regions of the conformational space, pointing towards the impact of induced fit in RNA-protein recognition.

## METHODS

### Atomic-resolution structural data and RNA equivalence classes

The primary source of RNA structural data is the Protein Data Bank (PDB). This database is biased towards those structures that are determined more easily experimentally and is highly redundant, which means that the mining of PDB is required to obtain a non-redundant set of structures (12). Leontis group classified types of redundancy and they generate weekly lists of “Equivalence Classes” of RNA in their website (“http://rna.bgsu.edu/rna3dhub/nrlist”). Each Equivalence Class contains all the RNA redundant chains and a representative structure. Equivalence Classes contain RNA chains rather than complete PDB IDs. A non-redundant set of structures is provided by the list of Equivalence Class Representatives. They offer different lists of X-Ray structures according to a resolution threshold, and an additional list called “All” including all experimental conditions. For our purpose, the “All” list was used here, as defined in the release version 3.134.

### Non-redundant datasets

From Leontis list, different sets of non-redundant structures were obtained: *complete-dataset* using all Equivalence classes, *protein-subset* when the RNA is in contact with a protein, and *naked-subset* which contains free RNA. In addition, the *complete-dataset* was also divided into three additional subsets representing the determination techniques: *xray-subset, nmr-subset*, and *em-subset* for X-ray, NMR and Cryo-EM techniques, respectively. For the X-ray and Cryo-EM structures, a resolution threshold of 2.4 Å was used to ensure a right description of the sugar puckering (13). We considered only RNA chains with 3 or more nucleotides, the minimum number to measure the eta-theta parameters. Next, all the nucleotides were analysed and divided in two sets according to their puckering state (North or South). North nucleotides were selected according to these criteria: delta = 84° ± 30°; phase between -18° and 54°; and Dp distance (base-phosphate perpendicular distance, defined in (13))>2.9 Å. South nucleotides were defined with: delta = 147° ± 30°; phase between 126° and 198°; and Dp distance <=2.9 Å. Nucleotides that lacked atoms or whose bond distances exceed the threshold of 2 Å were considered as “broken” and excluded. Nucleotides obtained using X-ray with the phosphate or the C4’/C1’ atom with b-factor over 60 were also excluded (8). The final nucleotides in the dataset were restricted to the A, U, G, and C residues that have a complete backbone starting from the sugar of the “i-1” nucleotide (5’ neighbour) to the sugar of the “i+1” (3’ neighbour).

### Definitions and calculations

Once the list of structures was compiled, we computed the geometrical analysis of the RNA backbone. η–θ and other dihedrals were defined as in Wadley’s work (7). Puckering was measured as in Westhof’s (14). Stacking was represented geometrically using the R vectors defined by Bottaro (15) and quantified using a QM approach (see below). The calculation of the distance between points in the η–θ map and the scoring of η–θ clusters were carried out as described in Wadley’s (7). RMSD, εRMSD (15), torsions and distances were computed in R using our package veriNA3d (11). RMSD calculations included backbone atoms from C4’_i-1_ to C4’_i+1_ or C1’_i-1_ to C1’_i+1_ in the case of η’–θ’. For the other nucleotide descriptors such as base pairing and η’’– θ’’ DSSR software was used (9).

### Ramachandran-like plots

The η–θ plot of North nucleotides revealed a large, highly populated region near its center (Figure 2). This region contains nucleotides with a backbone conformation typical of an A-RNA helix, so we refer to it as the *helical* region. It has a density that is almost two orders of magnitude higher than any other region of the η–θ map. To study the surrounding regions with precision, we separated the set of North nucleotides into *helical* and *non-helical* nucleotides like in Wadley’s work (7). Separate η–θ maps were so constructed for the *non-helical* North nucleotides and all South nucleotides. The same approach was applied to η’–θ’ and η’’–θ’’ plots. For each plot, the density was measured using a Kernel Density Estimation (KDE), with 361×361 grid points and a bandwidth = 40. From the densities at each point of the η–θ map, the mean (*ρ*) and standard deviation (*σ*) were calculated and contour maps at 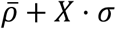 (using X = 1, 2, and 4) highlight High Density Regions (HDRs) or clusters.

**Figure 2.**
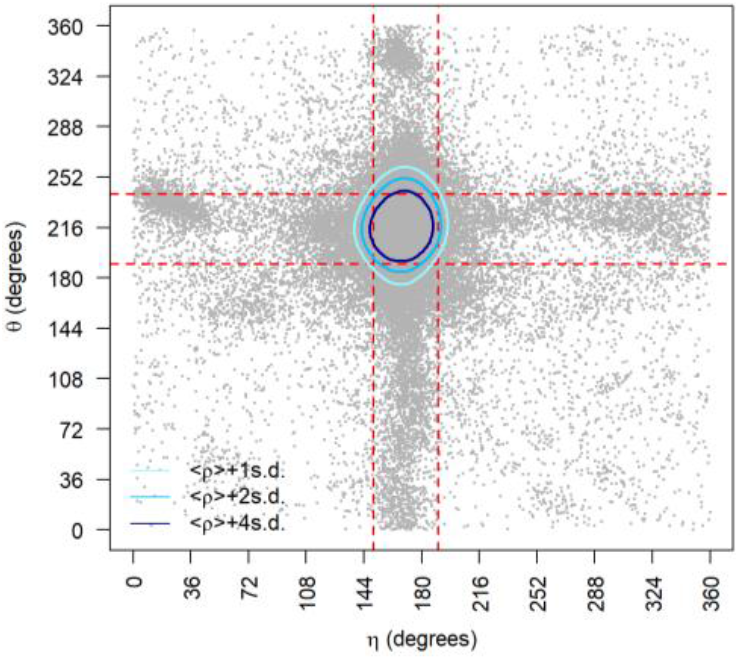
η-θ conformational space for all North nucleotides. Density contours of 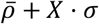 (X = 1, 2 or 4; light blue, blue and dark blue respectively) highlight a central cluster with a significant population of nucleotides, corresponding to the A-form helical conformation of the backbone. Red dashed lines delimit the region occupied by canonical helical A-form

### Stacking energy calculation

At the trinucleotide level, three combinations of stacking interactions were computed: the central base (i) with the nucleotide at its 5’-side (i-1), the central base with the nucleotide at its 3’-side (i+1), and (i-1)-to-(i+1) direct interaction that doesn’t involve the central base. To compute the stacking energy, the backbone of each trinucleotide was removed and replaced by a methyl group. Then the geometry of the individual bases was replaced by that optimized at the MP2/cc-pVTZ level of theory (16). Quantum mechanical energies were calculated at the dispersion-corrected (17, 18) DFT level B3LYP-D3(BJ) with the double-polarized def2-TZVPP basis set (19) and interaction energies were corrected by the basis-set superposition error (BSSE) following Boys and Bernadi counterpoise method (20).

## RESULTS AND DISCUSSION

The continuous growth in number of experimental RNA structures demands for periodic updates of the analyses based on structural data mining. Since the seminal work of Wadley et al. (7) in 2007, the number of deposited RNA structures in public databases like the PDB has increased by a factor of 4.5 (from 1,333 in 2007 to 5,978 in 2022) (Figure S1), providing a quite complete picture of the RNA conformational state. We consider here the entire structural space as well as subspaces separating them by the experimental technique used to describe the structure. The classification presented in Table 1 shows the scope of each method in describing the RNA conformational space. In the same way, the *complete-dataset* was filtered in “naked” (*naked-subset*) and “interacting” (*protein-subset*) RNA fragments with the purpose of addressing the possible effects on RNA folding introduced by interacting partners ubiquitously found in biological contexts. Statistics on the number of nucleotides and RNA structures analysed in this work are reported in Table 1. Finally, we analysed the sequence dependence on the η-θ space and explored how to relate transitions in η-θ clusters with the rotation of canonical backbone torsions α, β, γ, δ, ε, ζ and χ.

**Table 1.** Summary of datasets used in this work and number of nucleotides and structures included in each of them.

|  | Structures |  | Nucleotides |  |
| --- | --- | --- | --- | --- |
|  | Total | North | South | Total |
| Pyle dataset (7) | 73 | 6813 | 811 | 7624 |
| complete-dataset | 1,756 | 94,696 | 15,461 | 110,157 |
| xray-subset | 1,109 | 45,299 | 6,946 | 52,245 |
| nmr-subset | 601 | 12,194 | 1,471 | 13,665 |
| em-subset | 56 | 47,156 | 8,774 | 55,930 |
| naked-subset | 1,236 | 42,632 | 5,042 | 47,674 |
| protein-subset | 850 | 47,997 | 9,761 | 57,758 |

### How much did the global η-θ RNA pseudo-torsional space changed from its conception to date

The η-θ pseudo-torsion (Figure 1), was classified according to the sugar-puckering conformation (North non-helical and South) and analysed following the same strategy reported by Pyle and co-workers on our *complete-dataset*. Surprisingly, although we dramatically increased the number of analysed structures (Table 1) our results for η-θ showed roughly the original distribution, except for two unreported high-density regions (HDRs) for non-helical nucleotides with North sugars (herein clusters numbered VII and VIII in Figure 3A); and one new cluster in the η-θ/South space (cluster V, Figure 3B). These results reveal only moderate changes in the sampled experimental η-θ space, indicating that, either the RNA motifs space was already extensively explored by the X-ray structures available at that time, or that certain motifs remain elusive to experimental techniques.

**Figure 3.**
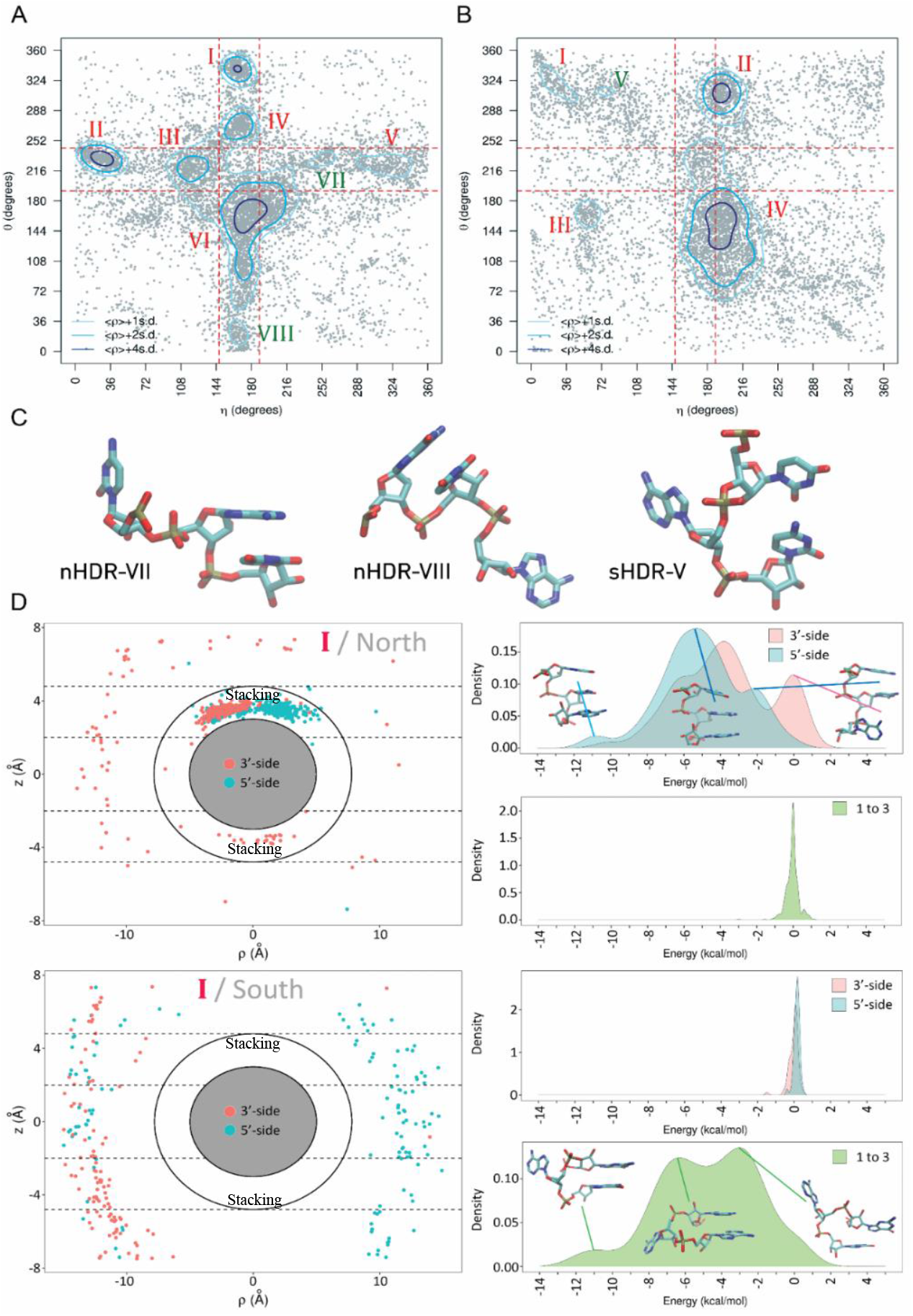
A) η-θ from the complete-dataset for non-helical nucleotides with sugar conformation in North. Density contours of 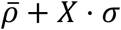 (X = 1, 2 or 4; light blue, blue and dark blue respectively) highlight regions of the plots with a significant population of nucleotides. Cluster or High Density Regions (HDRs) previously identified by Pyle and co-workers are labelled in red, while new clusters found in this work appear in green. Red dashed lines delimit the region occupied by canonical helical A-form. B) Same as (A) for nucleotides in South configuration. C) Representative structure for each new cluster found. D) Example of the stacking analysis for cluster I North (top) and cluster I South (bottom). The panel on the left shows the ρ and z components of the Rvector (15) between the given nucleobase and its 5’ (blue) and 3’ (red) neighbouring nucleobases. The inner space between ellipses and dotted lines labelled as “Stacking” shows the region in which stacking occurs. The panels on the right show the interaction energy between the given nucleobase and its 5’ (blue) and 3’ (red) neighbouring nucleobases (top panel) and the direct interaction energy between the 5’ and 3’ nucleobases (bottom panel).

The non-helical North HDRs described originally by Pyle and co-workers have either helical η (145° < η < 190°) or helical θ (190° < θ < 245°) values. Notably, the new clusters found in the η-θ North space (VII and VIII) have the same pattern. Therefore, all HDRs fall within the red dashed lines in Figure 3A, indicating that North nucleotides are usually located in the frontier of helical motifs. In contrast, the RNA nucleotides with sugar moieties in C2’-endo are, as expected, mostly found outside the helical region defined by red dashed lines in Figure 3B.

We investigated the structural composition of the identified clusters using the RMSD values for all trinucleotides of a given HDR, and a representative one was determined as the most similar to the others in the same region (the structure of all representative nucleotides is shown in Figure 2C for new clusters reported herein and Supp. Figure S2). Using Pyle’s similarity score (7), see Table 2; we found that the new clusters VII-VIII in η-θ/North and cluster V in η-θ/South displayed high similarity percentages (98.8, 96.0 and 93.3 respectively), and therefore they are formed by a group of nucleotides with the same backbone conformation.

**Table 2.** Summary of clusters’ similarities, and the number of PDBs in each cluster for η-θ conformational space.

| Puckering | Cluster | Similarity | n |
| --- | --- | --- | --- |
| North | I | 97.3 | 404 |
| North | II | 99.2 | 600 |
| North | III | 89.5 | 327 |
| North | IV | 95.2 | 309 |
| North | V | 93.8 | 242 |
| North | VI | 73.2 | 1,622 |
| North | VII | 98.8 | 81 |
| North | VIII | 96.0 | 124 |
| South | I | 87.0 | 138 |
| South | II | 94.0 | 631 |
| South | III | 93.2 | 148 |
| South | IV | 68.1 | 2,457 |
| South | V | 93.3 | 15 |

The stacking was described geometrically using the R-vector from base(i) to base(i±1) defined by Bussi group (15) and energetically from thousands of QM calculations (see Methods). In this way, a systematic description of each HDR for both South and North sugar conformations was obtained (see Table S1). Results in Figure 3D for clusters I in both North/South sugar conformations, display a range of energy that covers values from slightly repulsive stacking contributions to very stable ones (2 to -14 kcal/mol). Note how stacking interactions of the central nucleotide with its nearest neighbours are key for explaining conformations found in North cluster I (nHDR-I), while 1 to 3 interactions are the main players in stabilizing the structures found in cluster I in South (Figure 3 D).

The intramolecular hydrogen bond capability of the 2’OH group of the ribose in RNA (21), is another corner stone contact among HDRs found far from the helical region. As described in detail in Table S1, the 2’OH-phosphate interaction is a contact found prevalently in clusters I, II and V in η-θ/North. This shows the versatile role of this interaction in stabilizing non-canonical motifs while also serving as a primary molecular switch contributing to specific protein-RNA recognition (21). Our detailed analysis of each HDR and the correlations of η-θ pseudo torsions with all backbone angles also allowed for a more general observation: all high-density regions in the η-θ space are interconnected by a seemingly simple combination of torsional rotations (Figure 4). Transitions in the η axis appear mainly driven by changes in α (Figure 4A), while transitions in θ depend on changes in *ζ* (Figure 4B). This suggests an interpretation for RNA conformational transitions occurring at two levels: (i) a “coarse” level, that allows RNA to move between HDRs, and (ii) a “fine” level on how these movements translate to the all-atom space. This observation is developed below.

**Figure 4.**
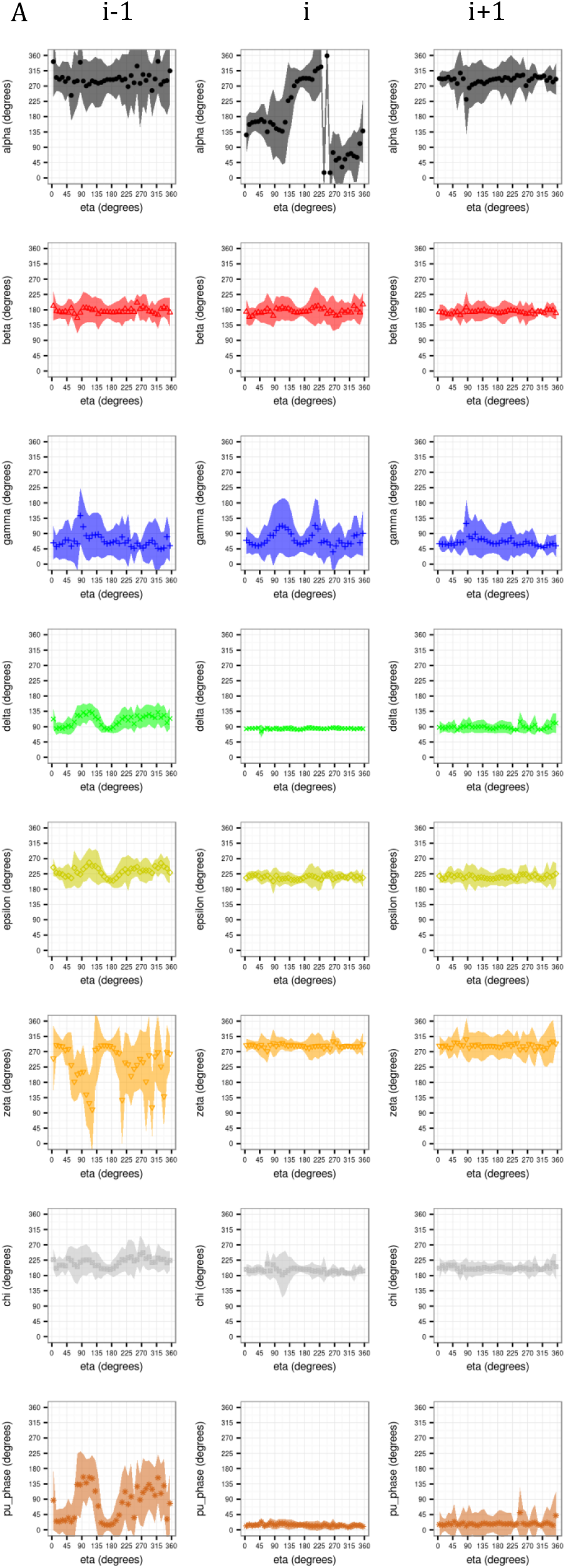
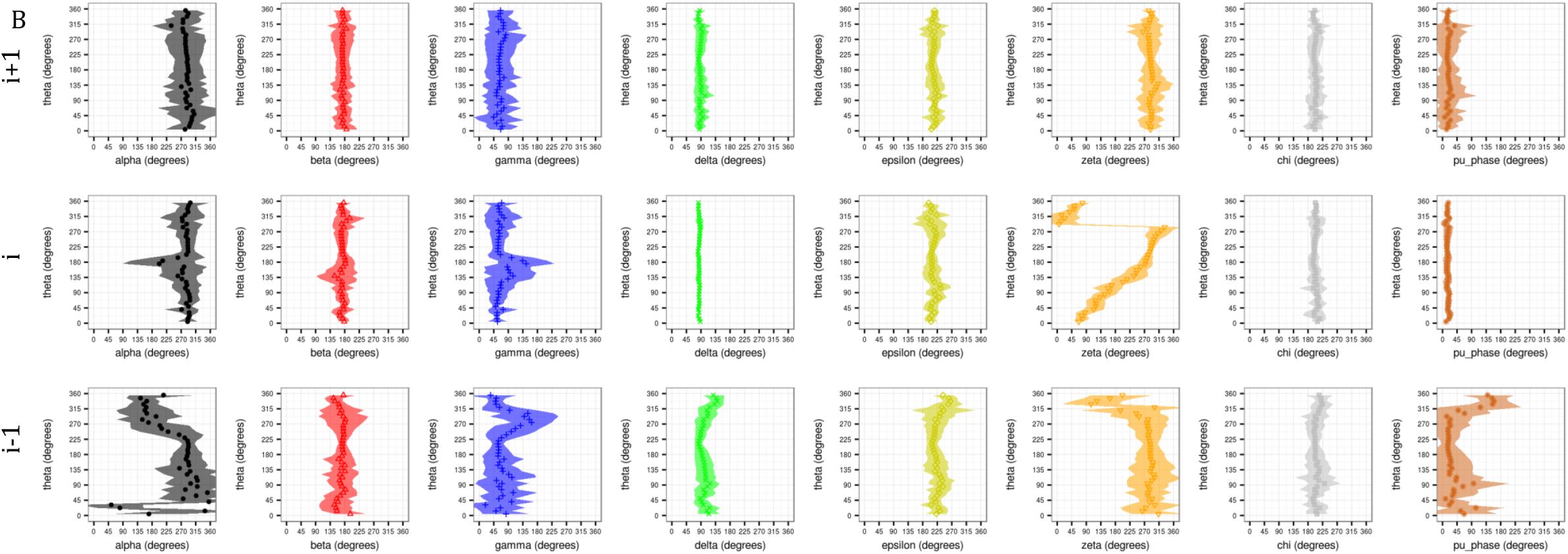
A) In previous page. Distributions of real dihedral angles in the nucleotides i-1 (panels on the left), i (mid-panels), and i+1 (panels on the right), as a function of the pseudo-dihedral angle eta (η) in the central nucleotide i. From top to bottom panels show: alpha (black), beta (red), gamma (blue), delta (green), epsilon (yellow), zeta (orange), chi (gray), and puckering phase (brown). B) Distribution of real dihedral angles in the nucleotides i-1 (bottom-panels), i (mid-panels), and i+1 (top-panels), as a function of the pseudo-dihedral angle theta (θ) in the central nucleotide i. From left to right panels show: alpha (black), beta (red), gamma (blue), delta (green), epsilon (yellow), zeta (orange), chi (gray), and puckering phase (brown)

### Global η’-θ’ and η’’-θ’’ RNA pseudo-torsional space

The pseudo-torsions η’-θ’ and η’’-θ’’ in their definition share similarities with the original η-θ torsions (Figure 1B,C). In η’-θ’ the C1’ atom is chosen instead of C4’, while in η’’-θ’’ the standard reference frame (SRF) as defined by Olson and co-workers (10) is used to account for the relative orientation of the base respect to the backbone. It should be noted that the location of the SRF depends upon the width of base-pair C1’-C1’ spacing and the pivoting of complementary bases in the base-pair plane, and hence to analyze single-stranded RNA an idealized base-pair reference is needed.

A first inspection of the η’-θ’ space (Figure 5A,B) revealed a very similar distribution to the one provided by η-θ, but with lower discriminating power as less clusters are visible in the North plane. Despite its reduce descriptive power, the η’-θ’ pseudo-torsional space might be advantageous for mapping X-ray models to their electron density maps, given that C1’ is used as a key reference atom to fit the experimental densities into molecular models (8, 22–24). Coarse-grained models for RNA (25) based on η’-θ’ could also benefit over η-θ since the C1’ atom is used to define the major/minor groove in curvilinear helicoidal coordinates (26), therefore the grooves’ dimensions would have a direct readout.

**Figure 5.**
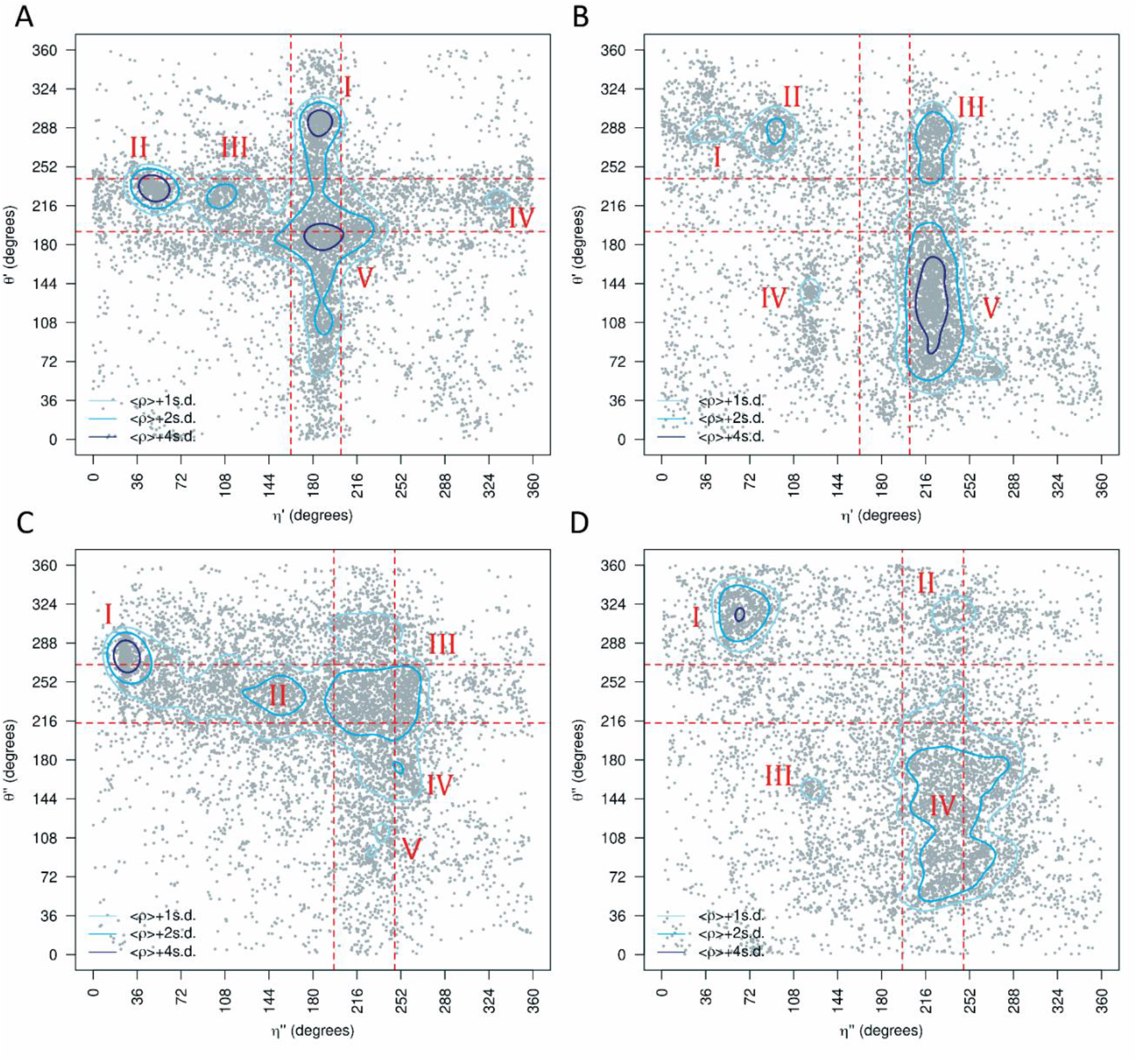
η’-θ’ and η’’-θ’’ conformational spaces for the complete-dataset. The results are divided into North non-helical (left plots) or South (right plots) sugar conformations. Density contours of 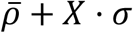 (X = 1, 2 or 4; light blue, blue and dark blue respectively) highlight regions of the plots with a significant population of nucleotides. Red dashed lines delimit the region occupied by canonical helical A-form (those structures are removed from the analyses as described in the Methods Section). Cluster numbers are identified in red. A) Densities clusterization of the η’-θ’ pseudo-torsions for central nucleotides in North. B) Same as (A) for central nucleotides in South. C) Densities clusterization of the η’’-θ’’ pseudo-torsions for central nucleotides in North. D) Same as (C) for central nucleotides in South.

To our knowledge, no data mining of PDB has been done using the η’’-θ’’ pseudo-torsional space. In our datasets, the η’’-θ’’ space looks similar to η-θ and η’-θ’ (Figure 5C,D) but the distribution is less detailed. As a result, the discriminating power is even lower than the η’-θ’ one, as North clusters I to IV belong to a same high-density region with contour 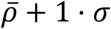. Each cluster is then a more heterogenic mixture of structures, and hence, η’’-θ’’ appears to be not so useful to describe and understand backbone conformational substate diversity as found experimentally.

In summary, even if the η’-θ’ definition might have some advantages in terms of modelling, the traditional η-θ space is preferable to describe the conformational space accessible to RNA.

### Experimental techniques describing RNA conformations

Originally, only high-resolution structures obtained by X-ray crystallography were used to analyse the η-θ space (7). However, to date, an important number of RNA structures were determined by means of solution NMR, and more recently by Cryo-EM which has reached a resolution close to that of X-ray while allowing for the determination of huge RNA complexes (27). We divided then the *complete-dataset* in 3 subsets, one for each method, in order to analyse their contributions and limits: *xray-subset, nmr-subset*, and *em-subset* (see Methods). As shown in Table 1, a significant number of nucleotides can still be analysed despite the division performed.

The *xray-subset* (Figure 6A,B), although being dramatically enlarged in the number of structures, reproduces well Pyle’s η-θ space (7). The *em-subset* (Figure 6C,D) and *nmr-subset* (Figure 6 E,F) show new clusters VII/VIII in the North plane and cluster V in the South plane. Clusters VII-VIII in η-θ/North are mostly exclusively populated by trinucleotides found in large ribosomal complexes that could have been determined with atomic resolution only recently (28). Apart from these two new clusters, trinucleotides in the *em-subset*, extracted from only 32 PDB structures (Table 1), reproduce by themselves all the conformational sub-states observed in the *complete-dataset*. Remarkably, ribosomes seem to exploit geometrically and energetically all the possible backbone sub-states to achieve their complex 3D folded structures that in turn depend on RNA-protein interactions. Interesting results are obtained when analysing the pseudo-torsional space revealed by solution NMR experiments (*nmr-subset*). As shown in Figure 6 E,F only certain conformations seem to be detected when using this technique (clusters II-III-IV-VI in North and IV-V in South), where large deviations from the canonical η-θ values are not frequently sampled. The new cluster V/South described herein for the *complete-dataset* (Figure 3B) is mainly composed by structures determined using NMR. Furthermore, focusing on the *nmr-subset*, the large and heterogeneous cluster IV/South can be divided into two clearly distinct populations (Figure 6F).

**Figure 6.**
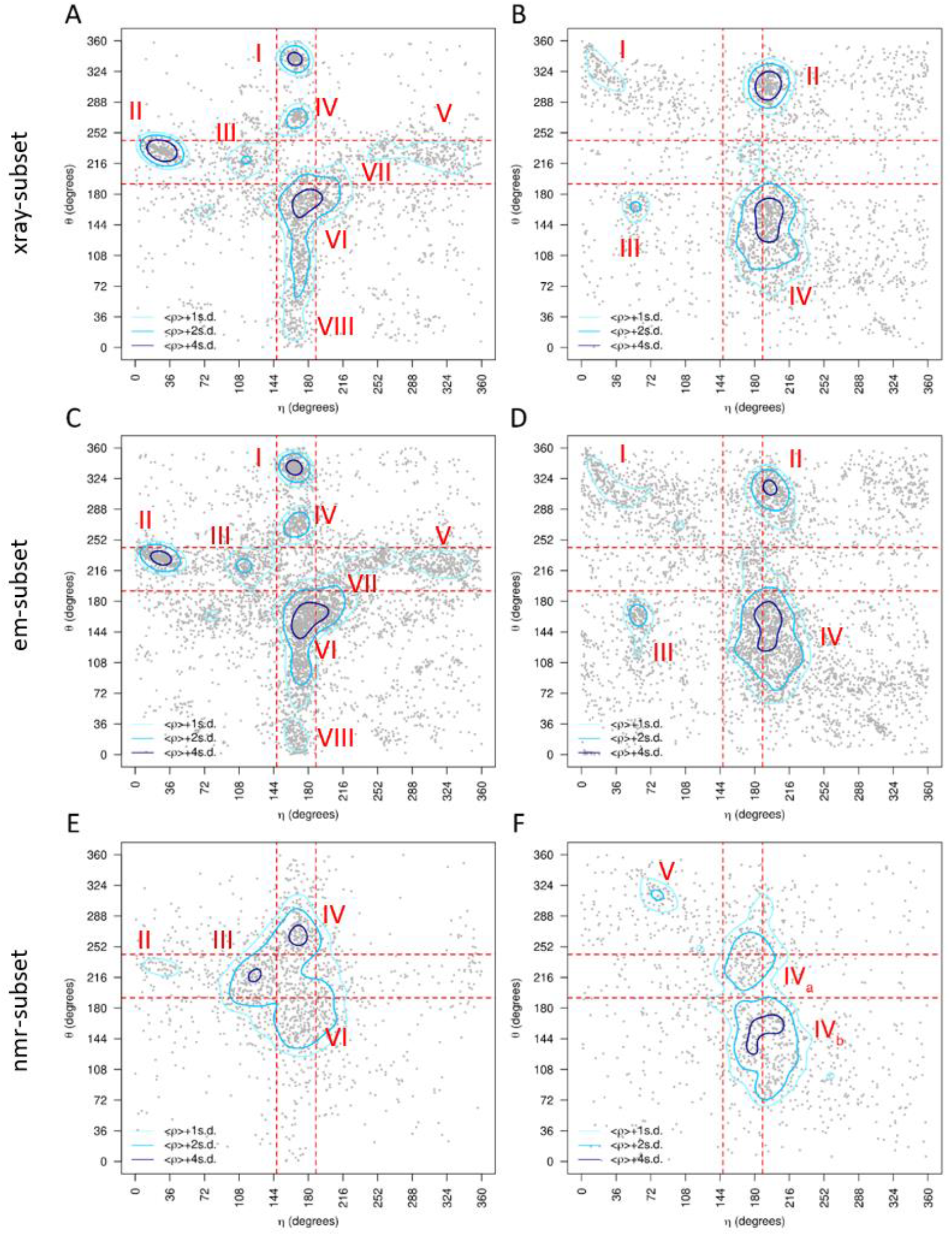
η-θ conformational space as detected by each experimental technique. Results are divided into North (panels on the left) or South (panels on the right) sugar conformations. Density contours of 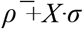 (X = 1, 2 or 4; light blue, blue and dark blue respectively) highlight regions of the plots with a significant population of nucleotides Red dashed lines delimit the region occupied by canonical helical A-form (those structures are removed from the analyses as described in the Methods Section). Cluster numbers are identified in red. A) Densities and cluster analysis of the xray-subset for central nucleotides with sugars in North. B) Same than (A) for South conformations. C) and D) are the same as (A) and (B) for the em-subset. E) and F) are the same as (A) and (B) for the nmr-subset.

Being based on completely different approaches and physical observables, which make comparisons rather difficult, the main differences among these 3 methods might arise from features like the experimental temperature at which data collection is produced. While CryoEM typically cools down the sample between 4 K (liquid helium cryogen) and 77 K (nitrogen, propane or ethane coolers) (29), the structures from the *xray-subset* were collected at 112 ± 44 K and NMR ones at near NTP conditions 293 ± 28 K. Above a given temperature threshold, thermal fluctuations of RNA molecules in solution might be preventing the transient stabilization of some extreme and less stable backbone sub-states. Consequentially some sub-stated detected by CryoEM and Xray may not be captured when performing NMR experiments. Other difference between these 3 methods might arise from the different size of the RNA molecules in each of the datasets. Apparently, this should not be relevant as the average chain length of the analysed structures is very similar between NMR and X-ray datasets (26 and 34 residues on average, respectively, while it is 323 for the *em-subset*). However, a closer look into our datasets indicates that both the X-ray and Cryo-EM subsets contain several long RNA chains such as ribosomes with thousands of nucleotides that can sample all the conformational space of the RNA backbone, whereas the longest structure in the NMR set is 155 residues (PDB ID: 2N1Q) that contains mostly A-form RNA. Apparently, the longer a RNA molecule is, the easier is to explore alternative non-helical conformations, but this phenomenon could also be influenced by the contacts with proteins, which is explored hereafter.

### How do proteins affect the RNA η-θ space

We divided the *complete-*dataset into “naked” RNA and protein-bound RNA. This allowed for the analysis of the intrinsic or “naked” conformations that RNA could adopt (*naked-subset*) and the effect produced by proteins (*protein-subset*). The η-θ plots for North puckering (Figure 7) reveal that, while the *protein-subset* (protein closer than 5 Å) roughly resembles the *complete-dataset*, the *naked-subset* shows a nucleotide distribution confined to the helical region neighbourhood (clusters III, IV and VI). Clusters I/III/IV show a noticeable reduction in their population, and no HDR are found near θ values of ∼0° (new cluster VIII in the *complete-dataset*). These differences are an evident consequence of the divergent η-θ density distributions in the absence and presence of proteins. That is, the density of the *naked-subset* biases the sampling of the *complete-dataset* towards near-helical regions, while proteins preferentially enrich conformations far from the helical centre. Summarizing, these results indicate that proteins can stabilize North pucker RNA segments with η and θ values far from the helical conformation, which are less sampled in “naked” RNA. In the case of South puckering nucleotides, the absence of proteins enriches cluster IV in the helical region, splitting the cluster in two sub-HDR regions.

**Figure 7.**
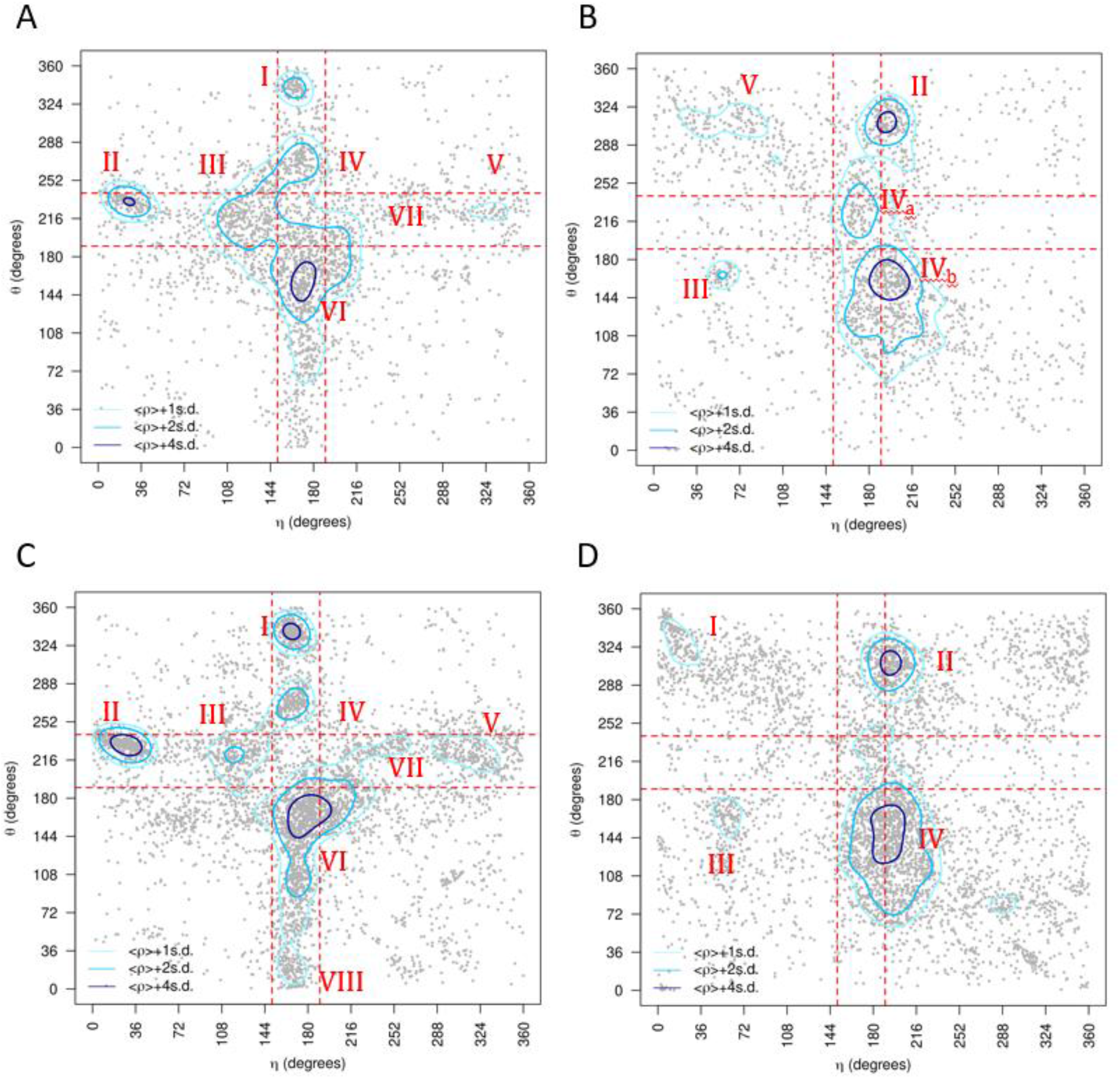
η-θ conformational space described by the naked-subset and protein-subset. Results are divided into non-helical North (panels on the left) or South (panels on the right) sugar conformations. Density contours of 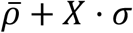 (X = 1, 2 or 4; light blue, blue and dark blue respectively) highlight regions of the plots with a significant population of nucleotides. Red dashed lines delimit the region occupied by canonical helical A-form (those structures are removed from the analyses as described in the Methods Section). Cluster numbers are identified in red. A) Densities and cluster analysis of the naked-subset for central nucleotides with sugars in North. B) Same as (A) for South conformations. C) and D) are the same as (A) and (B) for the protein-subset.

**Figure 8.**
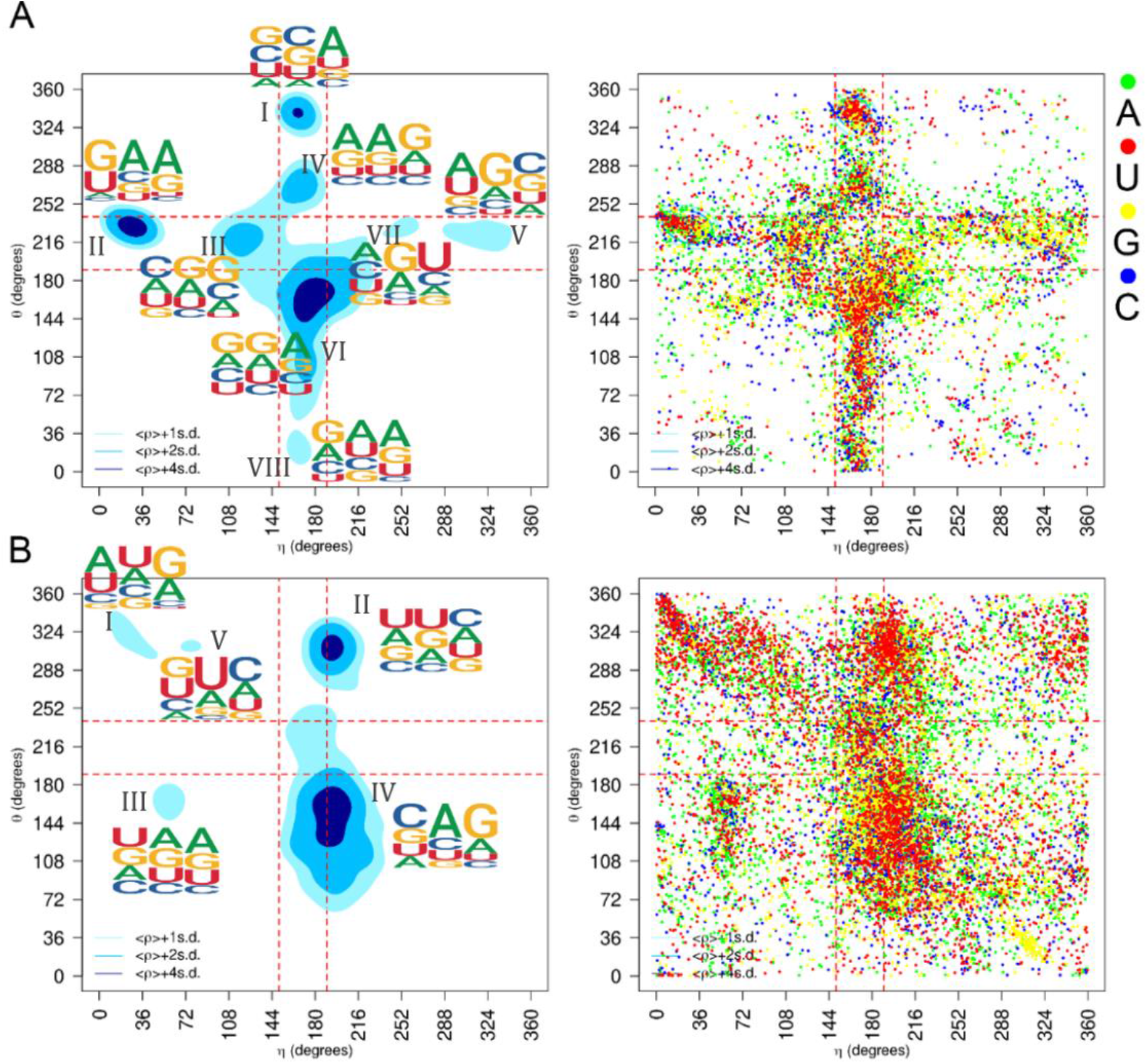
η-θ conformational space of the complete-dataset. Results are divided into non-helical North (top-panels) or South (bottom-panels) sugar conformations. Density contours of 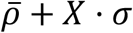 (X = 1, 2 or 4; light blue, blue and dark blue respectively) highlight regions of the plots with a significant population of nucleotides. Red dashed lines delimit the region occupied by canonical helical A-form (those structures are removed from the analyses as described in the Methods Section). Cluster numbers are identified in black in the left panels. A) The panel on the left shows densities and sequence logo-plots of the complete-dataset for sugars in North. The panel on the right shows the corresponding dot-plot with adenine (green), uracil (red), guanine (yellow) and cytosine (blue). B) Same as (A) for South conformations.

#### Sampling of the η-θ space is nucleobase dependent

The large number of nucleotides in each cluster allows for a reasonable analysis on the sequence dependencies in the η-θ space. Notable differences are observed when the *complete-dataset* η-θ map for North sugar rings is decomposed into nucleobase identity as evidenced in Figure 4.3.7. Despite fluctuations in their relative populations, clusters nearby the helical region (III, IV and VI), and cluster II are clearly observed for all four nucleobases, as well as South cluster II and III. However, in North, cluster I is depopulated of adenine, cluster V is almost exclusive of guanine, and uracil is almost absent in clusters V and VII. Considering that these later clusters are more populated in protein-RNA complexes (see previous sections), this result suggest that the nucleobase dependent sampling of the η-θ space is, in part, a consequence of sequence specific protein-RNA contacts. Furthermore, the presence of purines in the trinucleotide motif displayed as logo plots is highly dominant outside the helical region. In the case of South sugar rings, nucleobase dependence has been also observed, with cluster I almost exclusively populated by adenine and uracil, and adenine being the main responsible for cluster IV.

### A small set of torsional perturbations applied to a helical structure fully explain the RNA conformational diversity

All the North η-θ clusters fall into the space of helical η (∼160°) or helical θ (∼215°) (see the red lines in all our η-θ maps). Therefore, in most cases the backbone can move from one cluster to another just by rotating η or θ. For example, to go from the HDR-II (η=30°, θ=215°) to the HDR-III (η=110°, θ=215°), the backbone only needs to move along the η axis. To acquire insight on the specific torsion changes coupled to η and θ, average values of α, β, γ, d ε, ζ, χ and puckering were calculated for η and θ values in 5 degrees windows. This was done for the central, and for the first 5’-and 3’-neighbours. Figure 4 suggests that simple combinations of specific torsional perturbations encode the transitions between nHDRs. We have shown before that the torsions of η and θ can be traced by the movements of the real dihedral angles α, β, γ, d ε, ζ, χ, and puckering (Figure 4). Considering the η direction, discrete perturbations in ε, ζ, and puckering in the 5’-neighbour, and in α and γ in the central nucleotide result in the sequential transitions: nHDR-II↔nHDR-III↔helical↔nHDR-VII↔nHDR-V. Considering the θ direction, perturbations in α, γ and ζ for the central nucleotide, and in α, γ and puckering for the 3’-neighbour, explain the sequential transitions: nHDR-I↔nHDR-IV↔helical↔nHDR-VI↔nHDR-VIII. These results suggest that moving from the helical conformation to any of the nHDRs can be achieved by specific combinations of discrete torsional perturbations. This result implies that the complete North puckering RNA conformational space can be rationalized as a helical structure plus a small set of specific combinations of torsional perturbations.

## CONCLUSIONS

We explored here the conformational space of the “experimental” RNA backbone using our package veriNA3d (11). The large size of the database allowed to make subsets of the data and explore relationships that could have not been addressed before, such as the effect of proteins on the RNA backbone, or intrinsic differences due to the source of experimental structures. In addition, alternatives to η-θ to study the backbone conformation have been used, such as the η’-θ’ (which uses the C1’ instead of C4’) and the η’’-θ’’ (which uses a standardized point in the base plane instead of the C4’). We explored these alternative maps but found out that they tend to form less clusters and more heterogeneous, so we focused our analysis on the original η-θ space.

Quite surprisingly, the updated η-θ maps, show overall a similar general distribution to that found in Pyle’s 2007 study (7). However, we detect the presence of three small new clusters that had not been seen before and appear mostly due to the introduction of no X-ray techniques. In general, the X-ray and Cryo-EM subsets show almost equivalent η-θ maps, while the NMR subset is much more limited, covering only the clusters close to the helical A-form. One plausible explanation is that X-ray and Cryo-EM datasets include very large structures with thousands of nucleotides (e.g. ribosomes) and proteins that expand the conformational space that can be sampled by RNA in the non-helical η-θ space. Overall, it is clear that complementary sources of data are needed to fully recover the landscape of RNA backbone conformations.

Comparing samplings in *naked-subset* and *protein-subset*, we find evidence that protein contacts allow the RNA backbone to adopt the most extreme conformations. This agrees with previous findings in which proteins are shown to modulate the RNA conformation through contacts with the 2’OH (21).

We also find that the transitions between clusters can be traced by the rotation of specific dihedral angles, and only few changes are necessary to go from one cluster to another. Given that the η-θ maps are particularly interesting for coarse-grained applications in modelling RNA (25), our finding introduces a new way of moving in the η-θ space by using the real dihedral angles.

## Supporting information

EtaTheta_SuppInfo

