## Supplementary material for "The Pseudo-Torsional Space of RNA": EtaTheta_SuppInfo

### SUPPORTING TABLES

**Table S1**. Detailed description of all HDR derived from our *complete-dataset*.

| Cluster / HDR | Puckering | Description |
|---|---|---|
| I | North | This region has been reported to be structurally homogeneous, containing many nucleotides from S1 and S2 motifs, stacked between both 5'- and 3'- neighbours, the sugar ring of the latter frequently found in south. The update to the *complete-dataset* confirms the same observations for the dominating RMSD cluster, and also indicates that such conformation requires the central and first 5'-/3'- neighbours to form base pairs with surrounding RNA segments, although minor levels of unpaired states are also allowed. Regarding the type of pairing, the 5'- nucleotide forms mainly canonical cWW base pairs, the central nucleotide forms cWW, tHS or tSH base pairs, and the 3'- nucleotide forms tHH or tWW pairs. The most relevant contact observed in this HDR occurs between the O2' atom of neighbouring nucleotides and the phosphate oxygen atoms of the central nucleotide and the first 3' neighbour (probability(O2'-OP2)=0.79/0.68 and probability(O2'-OP1)=0.90/0.96, respectively. Worth noting, the latter interaction directly affects the α torsion of the 3' neighbour, which is one of the main torsional perturbations in nHDR-I. Finally, mild probability contacts are also observed between $Mg^{2+}$ cations or protein positive residues, and the 3' neighbour phosphate group. |
| II | North | This HDR corresponds to hairpin motifs with a well-defined nucleobase orientation signature, *i.e*. parallel stacking with the 3' neighbours, and loss of stacking with the 5' neighbours which are in anti-parallel orientation and located toward the same side of the 3' neighbours. In contrast to the other nHDRs far from the helical region, the contacts pattern of nHDR-II reveals that the unpaired base state is the most probable one for the central and 5'/3' nucleotides, non-sequential stacking has low probabilities and contacts with protein residues or $Mg^{2+}$ ions are negligible. On the other hand, high probability contacts are observed between the O2' of the 5' neighbour and the nucleobase of the 3' neighbor. In addition the phosphate group of the central nucleotide also shows high probability contacts with nucleobase polar atoms, although the orientation of the neighbouring nucleobase (that of the 5' neighbour) indicates such contact is a consequence rather than a cause of the local conformation. As a consequence, in the case of the n-HDRII, the most relevant interactions correspond to intramolecular contacts between nucleobase polar atoms and the O2' or phosphate group. This characteristic might underlie the persistence of this HDR regardless of the database decomposition (*naked-RNA-subset*, *protein-RNA-subset,* etc). |
| III | North | As previously indicated by Pyle and coworkers is frequently populated by nucleotides with a south sugar ring 5'- neighbour and that often occurs in 3' halves of adenosine platforms. This HDR is reported to be somewhat more heterogeneous than the nHDR-I and nHDR-II. The main particularity of this cluster is localized in the orientation of the 5' |

| | | |
|---|---|---|
| | | neighbour, explained by a change from canonical to g- and t/g+ values of ε and ζ, respectively, and the reported change in sugar pucker from north to south, when going from nHDR-II to nHDR-III. Noteworthy, both HDRs present the same non-canonical a distribution close to trans/g+ conformations. These observations suggest the hypothesis that nHDR-III is an intermediate state in the transition between the helical conformation and nHDR-II motifs (*e.g*. GNRA-like tetraloops) in which a north↔south puckering transition, entangled with changes in a, ε and ζ torsions, constitute the atomic level mechanism of the conformational change. |
| IV | North | This region has been previously characterized by noting a shift in the 3'-neighbour nucleobase that precludes stacking with the central nucleotide, and as being often found as the last nucleotide in GNRA tetra-loops. The pairing profiles features unpaired states and canonical cWW pairs for the three nucleotides. In addition, tHW is observed for the 5'-neighbour and tHS/tSH are observed for the central nucleotide. In addition, tHW is observed for the 5'-neighbour and tHS/tSH are observed for the central nucleotide. Regarding the torsion distributions, only α and γ for the 3'- neighbour sample non-canonical trans values, which appear responsible for the shift of such nucleotide away from the helical axis and the subsequent loss of stacking with the central nucleotide. |
| V | North | Original analysis of this region described it as lacking a distinctive structural motif, although a ~180° angle between nucleobase planes of the central and 5'- neighbour was frequently observed, as well as a south sugar ring of the latter. This cluster shows a central and 3'-neighbour nucleotides conformation rather similar to that of nHDR-II, but with a rotated 5'- neighbour, as expected for such region of the η-θ plot. This indicates that nHDR-V is highly populated by conformations similar to nHDR-II, with a similar rotation of the 5'- neighbour, but in the opposite direction. Torsion analysis reveal that only α and ζ from the central and 5'-neighbour, respectively, have non-canonical distributions, with the former sampling the g+ region while the latter is spread overs g+, trans and g- regions. The base pairing profile indicate a strong preference of the central and 3'- neighbour nucleotides for canonical cWW pairing and a minor contribution of unpaired states, while the 5'- neighbour is mainly unpaired with minor contributions of cWW or tWW pairing. |
| VI | North | Originally this region was reported to be the result a small base plane twist between the central and 3'- neighbour, facilitating sugar edge pairing of the former. This feature is characteristic of the cross-strand stack motif, which contains nucleotides from tandem purine-purine base pairs. This is the HDR with the higher heterogeneity reported, where as a general feature is that moving towards lower θ values requires a shift of ζ from g- to trans. In addition to such changes in ζ, distortions on α, β and γ, are also observed depending on the value of θ. Base pairing statistics indicate that the central and first neighbour nucleotides are mainly canonically paired with minor contributions from tHS, tSH, tHW or tWH pairing or unpaired structures. |

| | | |
|---|---|---|
| VII | North | This cluster was not reported before. It is a HDR which fall down in typical θ-value helical region, but the η-value for nucleotides in this cluster is ~ 60° from helical. These residues are frequently succeeded on 3'-side by nucleotides with C3'-endo sugar pucker. |
| VIII | North | This original cluster was not reported before either. The notably region proximity with HDR VI suggests similar structural features . The η-value for a nucleotide in this cluster is typical for a helical nucleotide, but the θ-value for nucleotides in this cluster is ~160° from helical. These residues are almost always preceded on the 5'-side by nucleotides with C3'-endo sugar pucker. |
| I | South | This region was originally linked to specific motifs (kink- and p-turns), and are characterized by a splayed out nucleobase required to accommodate a sharp a kink. As previously reported by Pyle and coworkers, sHDR-I was split in two regions, one with η~0 and the other with η~72 (now described herein as new independent HDR labelled cluster V). The former (now sHDR-I) coincides with that previously observed by Wadley *et al*, showing the same structural characteristics. |
| II | South | Nucleotides in this region are often found as constituents of asymmetric internal loops *i.e.* second residue of a two nucleotide extruded helical strand. ). The main torsion perturbation from canonical values (besides the central nucleotide south puckering), is localized on ε of the central nucleotide. In fact, an ~60° rotation of only such torsion transforms sHDR-II representative structure into a canonical helical conformation, except for the central nucleotide ring being in south. In the absence of proteins (*naked-RNA-dataset*), essentially the same features are observed for sHDR-II, with the only difference being an increase of cWW pairing at the expense of unpaired bases for the central and 5'-neighbour. Overall, sHDR-II is revealed as a small perturbation on ε and sugar pucker from a canonical helical conformation, which, in the absence of proteins presents higher canonical Watson-Crick pairing, while proteins probably stabilize unpaired states. |
| III | South | Nucleotides in this region have been reported to be constituents of S1/S2 motifs. Analysis of datasets (protein, complete and naked) is consistent with this view, and shows a base pairing trend where the 5' neighbour is unpaired or engage in cWW or tSH/tHS pairing, the central base is unpaired or engage in tWW or tHH pairing, and the 3' neighbour is unpaired or engage in cWW pairing. Torsion distributions reveal non-canonical values, namely, the 5'- neighbour has ζ in the g+ region, the central nucleotide has α in trans, ε in g- and ζ in tans, and the 3' neighbour α, γ and ζ sample several sub-states while χ is in high anti. |
| IV | South | This region comprises nucleotides with heterogeneous motif identities (*e.g*. single nucleotide bulges, A-platforms, p-turns, and W-turns), with 5'-/3'- neighbours of varying sugar puckers. In agreement with this, higher heterogeneity in the RMSD values were observed compared to the other HDRs in the south η-θ space. |
| V | South | This original cluster not reported before appears as split of HDR I, and as was noted by Pyle the sharply non-helical η and θ values of nucleotides in this cluster is characterized by a splayed out nucleobase required to accommodate a sharp to a kink. The HDR I and probably this region is linked to the specific motifs kink- and p-turns. These residues |

are always preceded on the 5’-side by nucleotides with C3’-endo sugar pucker.

### SUPPORTING FIGURES

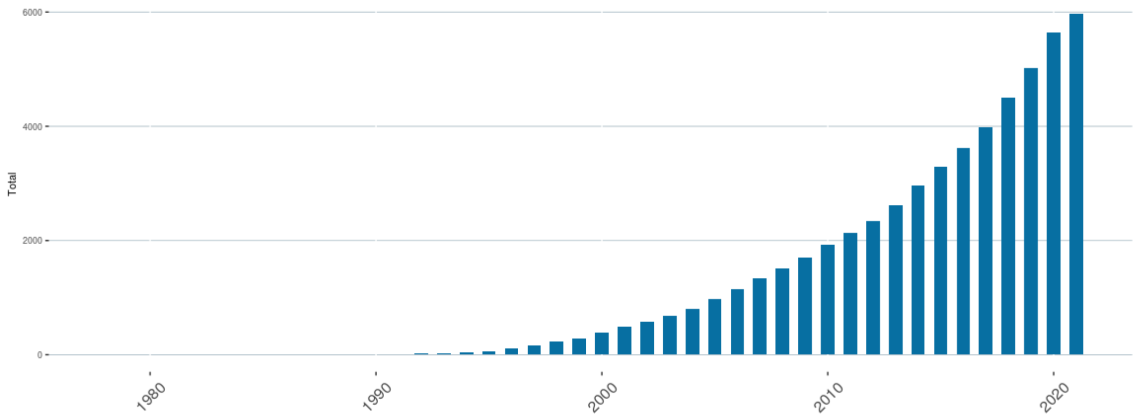


**Figure S1.** Evolution of the number of structures containing DNA deposited in the Protein Data Bank.

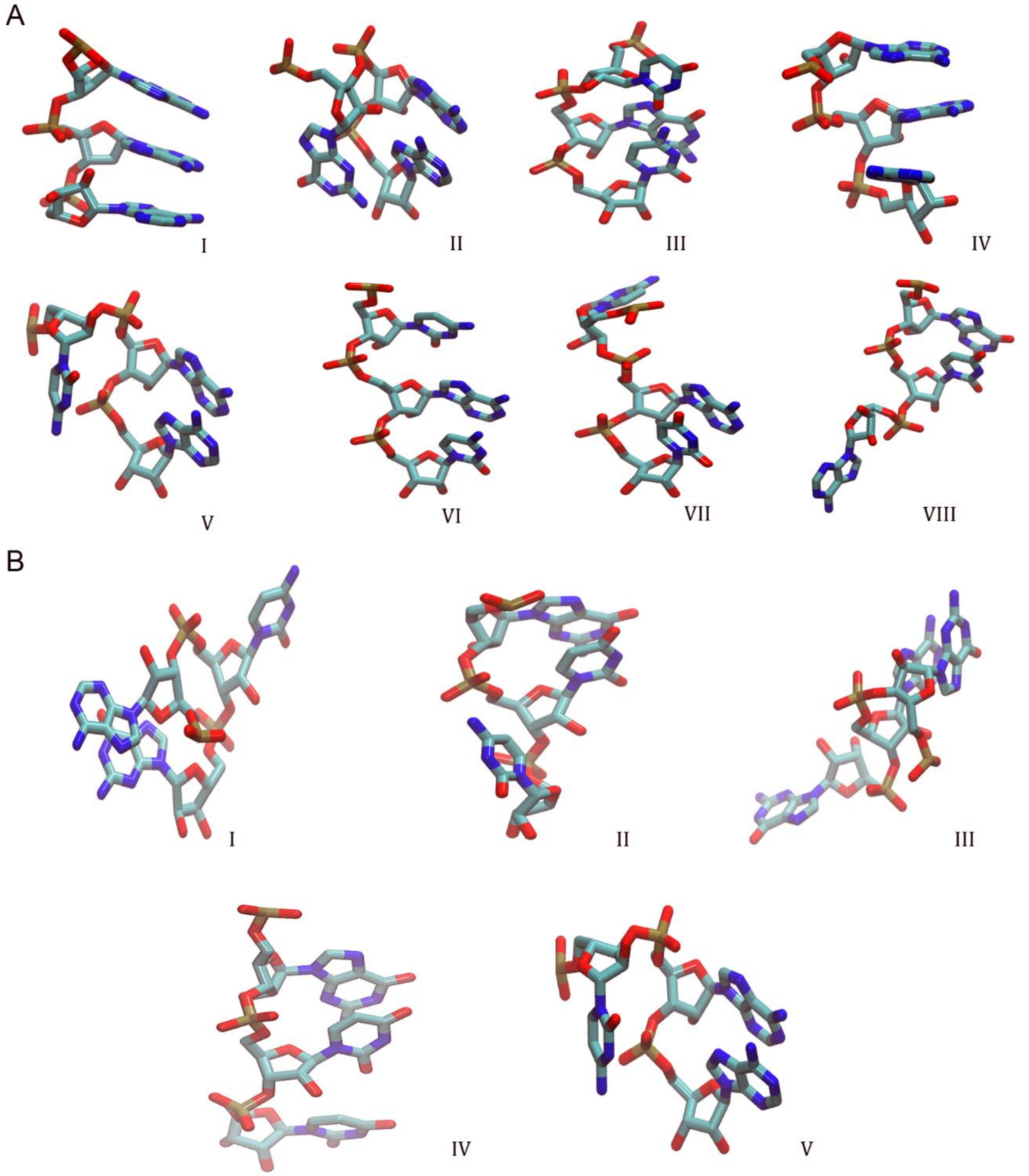


**Figure S2.** A) Representative structures for the eight clusters found in North conformation: Cluster I (PDBID 5J7L); Cluster II (PDBID 5J7L); Cluster III (PDBID 6AJ3); Cluster IV (PDBID 6SGC); Cluster V (PDBID 4Y4O); Cluster VI (PDBID 6SPB); Cluster VII (PDBID 5T5H); and Cluster VIII (PDBID 6V3A). B) Representative structures for the five clusters found in South conformation: Cluster I (PDBID 4V9F); Cluster II (PDBID 1F7U); Cluster III (PDBID 6V3A); Cluster IV (PDBID 4JF2); and Cluster V (PDBID 4V88).
